# Ribosomal s6 kinase (RSK) is a mediator of aquaporin 2 S256 phosphorylation and membrane accumulation after EGFR inhibition with erlotinib

**DOI:** 10.1101/2022.12.06.519384

**Authors:** Richard Samuel Edward Babicz, Pui W. Cheung, Noah Baylor, Richard Bouley, Dennis Brown

## Abstract

Vasopressin (VP) activates PKA, resulting in phosphorylation and membrane accumulation of aquaporin-2 (AQP2). Epidermal growth factor receptor (EGFR) inhibition with erlotinib also induces AQP2 membrane trafficking with a phosphorylation pattern similar to VP, but without increasing PKA activity. Here, we identify the ribosomal s6 kinase (RSK) as the final mediator phosphorylating AQP2 in this novel, erlotinib-induced pathway. We found that RSK was expressed in medullary principal cells in rat kidneys. RSK inhibition with BI-D1870 or siRNA blocked erlotinib-induced AQP2 S256 phosphorylation and membrane trafficking. CRISPR-generated RSK knockout cells failed to show increased S256 phosphorylation in response to erlotinib. Like PKA, RSK was able to phosphorylate AQP2 S256 *in vitro*. Stimulation of PDK1, a known activator of RSK, caused AQP2 S256 phosphorylation and membrane accumulation similar to erlotinib and VP. We conclude that RSK is the terminal kinase phosphorylating AQP2 at S256 upon EGFR inhibition by erlotinib.

## Introduction

A chief regulator of water balance in the body, vasopressin (VP) maintains water homeostasis and allows for adaptation and survival of organisms in variable climates. In response to decreased circulating blood volume or an increase in osmolality, the posterior pituitary gland releases VP into the circulation. VP then binds to the vasopressin receptor type 2 (V2R) on the basolateral membrane of principal cells in kidney collecting ducts and induces AQP2 apical membrane accumulation resulting in water reabsorption into the circulation to replenish blood volume.

The effect of VP occurs via the activation of the canonical signaling pathway centering around protein kinase A (PKA). When V2R is activated by VP, it stimulates adenylyl cyclase, which results in an increase in intracellular cyclic adenosine monophosphate (cAMP). cAMP activates PKA which induces AQP2 phosphorylation and accumulation in the apical plasma membrane [1–4].

Current therapies for patients suffering from dysregulation of water homeostasis target this canonical V2R-PKA-AQP2 pathway. In patients with excessive water reabsorption due to aberrantly increased VP production and activation of V2R receptor, a class of medications known as “vaptans” selective block the V2R [5]. While trials have shown some promise in populations with heart failure, cirrhosis, polycystic kidney disease, and SIADH [6–10], several drawbacks have limited clinical use of these drugs. Adverse effects include liver toxicity, rapid overcorrection of sodium, and progression of renal disease [11–13]. For diseases characterized by a deficit of water reabsorption, available pharmacotherapy also depends on modulation of the canonical V2R-PKA-AQP2 pathway. Nephrogenic diabetes insipidus (NDI) is characterized by excessively dilute urine resulting from an inability to traffic AQP2 to the membrane of the collecting duct [14]. Management of NDI centers around adequate access to drinking water, limiting solute intake, and drug therapy (eg. Desmopressin, NSAIDs) which upregulate the canonical VP pathway [15]. Unfortunately, these strategies often lack efficacy, and quality of life can suffer with some patients having to drink up to 20L of water daily [16]. The discovery of an independent pathway regulating AQP2 holds the promise of identifying an alternative therapeutic target to treat a wide spectrum of water balance disorders.

In recent years, several lines of investigation have shown that AQP2 trafficking may be affected by pathways operating independently of the known V2R-cAMP-PKA pathway [2]. Our group has shown that erlotinib, an EGFR inhibitor, increased AQP2 membrane accumulation without affecting the canonical cAMP-PKA signaling pathways [17]. Physiologically, erlotinib was able to decrease urine output and increase urine osmolarity in mice with lithium-induced NDI and rescued the polyuric phenotype [17]. EGFR activation attenuated AQP2 phosphorylation [17], and prior studies have shown EGF infusion in sheep and rats induces a water diuresis [18–20]. These findings suggest that EGFR signaling tonically inhibits VP signaling in water regulation. Following this line of inquiry we found that inhibition of c-Src, a downstream kinase of EGFR, also induced AQP2 membrane accumulation [21]. Clinical observations corroborate these findings, as patients on the same c-Src inhibitor used in our study (dasatinib) have increased prevalence of hyponatremia, or low blood sodium caused by excessive reabsorption of water from the kidneys [22]. These encouraging data suggest that EGFR and its downstream effectors have a potential role in water regulation, and prompted us to further investigate and dissect the pertinent EGFR pathways in kidney water balance.

The epithelial growth factor receptor (EGFR), also known as ErbB1, belongs to the ErbB family of receptor tyrosine kinases. While epidermal growth factor (EGF) is the principal ligand of the EGFR, up to six other lower-affinity ligands have been shown to bind the EGFR with varying effects in the kidney [23]. Activation of EGFR by EGF is associated with numerous pathways including the RAS-RAF-MEK-ERK MAPK pathway, the PI3K-AKT-mTOR pathway, phospholipase C-gamma PKC signaling, Cbl-mediated endocytosis, and Nck/PAK signaling [24, 25]. The RAS-RAF-MEK-ERK cascade is considered among the most important of these. In this pathway, EGFR ligand binding results in receptor autophosphorylation and activation of RAS and RAF. RAF in turn activates MEK1/2 which activates ERK1/2. One of ERK’s many targets is the ribosomal s6 kinase (RSK). Once activated by ERK, RSK translocates to the nucleus where it triggers cell growth via CREB phosphorylation as well as activation of c-Fos and cyclin D1 [25].

Our findings published here show that RSK is the likely final mediator in the novel pathway of AQP2 trafficking induced by erlotinib. That RSK and PKA share common substrates has been long noted in the literature [26]. RSK’s N-terminal kinase domain (NTKD) is genetically related to PKA, both being grouped in the AGC family of kinases [27, 28]. As such, RSK and PKA each possess two distinct tertiary structural motifs characteristic of this kinase family: an “activation loop” and a “hydrophobic motif.” For RSK and PKA alike, efficient phosphorylation of substrates depends upon a very specific interaction between these two motifs, with both enzymes sharing a predilection for substrates with arginine in the −3 position [27, 28]. Homology analysis of amino acid sequences of AQP2 c-terminus indicated that RSK could potentially phosphorylate S256. This, together with knowledge of RSK as an established mediator in the EGFR-MEK/ERK pathway, prompted us to further investigate the role of RSK in AQP2 signaling.

## Materials and Methods

### Reagents

All reagents were purchased from Fisher Scientific (Waltham, MA), if not specifically cited. The erlotinib, an EGFR inhibitor was purchased from Cayman (Ann Arbor, MI). The BI-D1870, inhibitor of the ribosomal S6 kinase (RSK) was purchased from Selleckchem.

#### Cell culture and treatment

The LLC-PK1 cells expressing c-myc–tagged AQP2 were grown in DMEM with 10% FBS and split two times a week as previously described [29–32]. No mycoplasma was detected on routine DAPI staining with Vectashield/Dapi (Vector Laboratories, Burlingame, CA). Cells were cultured to 90% confluency on either 35 mm Petri dishes or on 15 × 15 mm glass coverslips #1.5 (Ted Pella Inc., Redding, CA). They were starved two hours in DMEM medium without serum before treatment. Drugs were diluted in DMSO, and added directly to the cell medium. Therefore, the untreated cells received the same volume of DMSO as a negative control. VP (10 nM) was added for 15 min as a positive control. Cells were pre-treated 30 min with ribosomal S6 kinase (RSK) inhibitors (BI-D1870, 10 μM) before the addition of erlotinib (2 μM), the EGFR inhibitor. After 30 min of treatment with erlotinib, cells were fixed or lysed before being used for immunocytochemistry or western blot analysis as described below.

#### Immunocytochemistry

The immunofluorescence assays were performed using LLC-AQP2 cells as previously described [29, 33, 34]. After treatment, the treated cells were fixed 20 min in phosphate buffered saline containing 4% paraformaldehyde and 5% sucrose. After 3 washes in PBS, cells were permeabilized with Triton X-100 (0.1% in PBS, Sigma-Aldrich). Non-specific signal was blocked by 10 minutes incubation in PBS, 1% BSA solution. AQP2 was detected using undiluted culture medium containing mouse anti-c-myc IgG produced by the hybridoma cells (MYC 19E10.2 [9E10] ATCC® CRL1729TM) followed by a secondary anti–mouse IgG conjugated to Alexa-488 (10 *μ*g/ml; Invitrogen). Images were acquired on a Zeiss LSM800 confocal microscope with Airyscan detector (Apochromat 63x, N.A.=1.4, Carl Zeiss Microscopy LLC, White Plain, NY). All images were processed using Adobe Photoshop CS5 software (Adobe, San Jose, CA).

#### Kidney immunohistochemistry

All animal experiments were approved by the Massachusetts General Hospital Institutional Committee on Research Animal Care (Animal Protocol No. 2016N000040). Adult male Sprague-Dawley rats (Charles River Laboratories, Wilmington, MA) were housed individually and maintained in a temperature-controlled room regulated on a 12-hour light/dark cycle with free access to water and food. Rats were fed with normal chow (Pro-Lab Isopro MPH 3000, LabDiet, St Louis, MO).

Immunostaining of RSK was performed on rat kidneys. In brief, rats were anesthetized with 2% isofluorane inhalation. They were perfused through the left ventricle with warmed 37ºC PBS until kidneys were cleared of blood then the kidney tissues were perfused with periodate-lysine paraformaldehyde (PLP) fixative solution at the rate of 42 ml/min for 5 min. After perfusion, the kidneys were harvested, cut into approx. 1 mm thick transverse slices, and immersed in periodate-lysine paraformaldehyde (PLP) fixative overnight at 4ºC. After fixation, the tissues were rinsed five times in PBS, cryoprotected overnight in PBS containing 30% sucrose, and then covered with OCT compound tissue plus (Fisher scientific, Waltham, MA). Cryosections (5 *μ*m thick) were cut using a Leica 3050S Microtome (Leica Microsystems, Buffalo Grove, IL). Sections were picked up on positively charged slides (Thomas scientific, Swedesboro, NJ). After rehydration in PBS for 20 minutes, sections were incubated with 1% SDS in PBS for five minutes as an antigen retrieval step [35] followed by washing three times five minutes in PBS. Non-specific staining was blocked by incubation of the sections in Background Buster (Innovex Biosciences, Richmond, CA) for five minutes followed by incubation in PBS containing 1% BSA for 10 minutes. Sections were incubated with rabbit anti–RSK antibody (1 *μ*g/ml, Cell Signaling) overnight, washed 3 times 5 minutes in PBS, then incubated with donkey anti–rabbit IgG conjugated to Alexa-488 (6 *μ*g/ml, Jackson ImmunoResearch Laboratories, West Grove, PA) for 1 hour at room temperature and finally rinsed three times in PBS. Sections were mounted in Vectashield (Vector Laboratories, Burlingame, CA) and images were acquired using Zeiss LSM800 confocal microscope with Airyscan detector (Apochromat 63x, N.A.=1.4).

#### In situ kidney slice preparation for immunohistochemistry

The in vitro kidney slice model has been described previously [36]. In brief, rats were anesthetized and the kidneys were perfused with PBS as described above. After perfusion, the kidneys were harvested and cut into 0.5 mm slices using a Stadie-Riggs microtome (Thomas Scientific, Swedesboro, NJ). Before treatment, all kidney slices were equilibrated 30 minutes in CO_2_ saturated Hanks’ balanced salt solution (HBSS: NaCl 110 *mM*, KCl 5 *mM*, MgSO_4_ 1.2 *mM*, CaCl_2_ 1.8 *mM*, NaOAc 4 *mM*, C_6_H_7_NaO_7_ 1 *mM*, D-glucose 6 *mM*, L-alanine 6 *mM*, NaH_2_PO_4_ 1 *mM*, Na_2_HPO_4_ 3 *mM* and NaHCO_3_ 25 *mM*). The RSK inhibitor BI-D1870 (10 *μM*) diluted in DMSO or DMSO alone was added to the kidney tissues. After 30 minutes incubation, the negative control (HBSS) or the positive control (VP, 10 nM) solutions were added to the kidney slices. After 10 min incubation, the kidney tissues were immersed in periodate-lysine paraformaldehyde (PLP) fixative overnight at 4ºC. After rinsing the tissues five times, they were incubated overnight in PBS containing 30% sucrose and then covered with OCT compound. Cryosections (5 *μ*m thick) were cut as described above and stored at 4°C before immunofluorescence staining. Tissue rehydration and non-specific staining blocking were performed as described above. However, the sections were incubated with rabbit anti–AQP2 antibody (1 *μ*g/ml, Alomone Labs, Jerusalem, Israel) overnight, and subsequent incubation with donkey anti–rabbit IgG conjugated to Alexa-488 (6 *μ*g/ml, Jackson ImmunoResearch Laboratories, West Grove, PA) for 1 hour at room temperature. Sections were mounted in Vectashield (Vector Laboratories, Burlingame, CA) and images were acquired using Zeiss LSM800 confocal microscope with Airyscan detector (Apochromat 63x, N.A.=1.4).

#### Western blotting

Western blotting was performed as previously described. Briefly, LLC-AQP2 cells were grown to 90% confluency. Cells were incubated with drugs or DMSO for 60 min, followed by 15 minutes with VP/FK. After incubation, the cells were lysed in RIPA buffer supplemented with a Complete protease inhibitor cocktail (Sigma-Aldrich), EDTA (5 *mM*) and phosphatase inhibitors NaF (1 *mM*), and sodium orthovanadate (1 *mM*). Protein concentrations were determined using the BCA kit according to the manufacturer (Pierce, Rockford, IL). NuPage SDS sample buffer containing the solubilized proteins (20 *μg)* were incubated at 75°C for 10 minutes. Samples were run on a NuPage 4%–12% Bis-Tris Gel (Invitrogen) and transferred on to Immun-blot PVDF membranes (Bio-Rad, Hercules, CA). The membranes were blocked in PBS Tween-20, 0.1% (PBS-T) containing 5% nonfat milk for 1 hour at room temperature. The membranes were incubated overnight with primary antibodies against rabbit anti-AQP2, and an antiphospho-p256 AQP2 antibody (1:1000, Abcam). The membranes were then washed five times in PBS-T before a second antibody incubation with horseradish-peroxidase conjugated donkey anti-rabbit IgG antibodies (0.08 μg/ml, Jackson ImmunoResearch Laboratories). The enhanced chemiluminescence signal was acquired using a G-box mini imaging system and GeneSys image acquisition software (Syngene, Frederick, MD, USA). After visualization, membranes were stripped using a stripping buffer (Boston Bioproducts, Inc., Ashland, MA) for 30 minutes, and re-incubated with different primary antibodies as listed previously. Intensities of phosphoproteins were analyzed by Image J software and intensities were corrected according to the amount of total AQP2 on the same membrane. The total and the phosphorylated RSK expression in treated LLC-AQP2 cells were analyzed by western blot analysis as described above.

### In vitro phosphorylation assay

We tested the phosphorylation of AQP2 by RSK, or PKA as a positive control, in vitro. 1.15 μg of purified AQP2 or AQP2 S256A COOH-tail peptide were mixed with either RSK (16 nmol) or PKA catalytic subunit (5.6 nmol) (Promega, Madison, WI) in a total volume of 12 μl reaction buffer containing 40 mM Tris pH 7.5, 20 mM MgCl2, 0.1 mg/ml BSA, 50 uM DTT, 10 μM ATP, 10 μCi[γ-32P] ATP, and incubated at 30°C for 5 minutes. The reaction was stopped by boiling the samples in SDS-PAGE sample buffer. Samples were resolved on a 12% NuPage gel. The gels were dried using the DryEase mini Gel Drying systems (Invitrogen) according to the manufacture protocol. Protein phosphorylation was visualized using autoradiography (Hyblot ES, Thomas Scientific, Swedesboro, NJ).

#### siRNA methods

As previously described above, LLC-AQP2 cells were cultured to 50% confluency and transfected with 20 nM siRNA targeting RSK1 (Invitrogen, Charlestown, MA, USA) 5’-GCCACGAAAGCAUGGUUUAUCUAAA-3’ (RSK siRNA1) or control CT scrambled siRNA 5’-GCCAAGCGAGUAUUGUAUUCACAAA-3’ for 48 h using Lipofectamine RNAi/MAX according to the manufacturer’s protocol. After transfection and appropriate treatments, cells were lysed for RNA extraction or Western blot analysis by methods described above, or fixed and stained for immunocytochemistry as described above.

#### CRISPR knockouts

LLC-PK1 cells expressing AQP2 constructs were transfected with pCMV-cas9-OFP plasmids containing gRNAs specific for the *rps6ka1* gene. Plasmids were constructed using Geneart kit (Invitrogen) and guide RNA sequence 5’-TCAAAGTAGAAGGTATCAT-3’. Plasmids were transfected with lipofectamine 3000 (Invitrogen) according to the manufacturer’s instructions. Orange fluorescent protein (OFP) expressing cells were sorted into 96 well plates (∼1 cell per well) using a FACS Aria II cell sorting machine at the CRM Flow Cytometry Core Facility at Mass General Hospital. RSK expression was assessed by western blotting. Sanger sequencing showed a one base pair insertion at the CRISPR target site. Forward primer: 5’-AGTGCAGACTGGTGGTCCTA-3’ Reverse primer: 5’-GCCGTCATCCTCCATTAGGC-3’. Knockouts were confirmed by western blotting.

#### Data analysis

Data are expressed as mean ± SD, and statistical analyses were performed as appropriate using the one-way Anova with multiple comparisons tests with Tukey correction. T-test was used when applicable. Differences were considered to be significant at P-value < 0.05. Specific P values are given in the figures.

## Results

### Inhibition of ribosomal-s6-kinase (RSK) decreases erlotinib-induced phosphorylation of AQP2 at serine-256 (S256)

In our prior study, we found that EGFR inhibition with erlotinib induced a pattern of AQP2 phosphorylation that was similar to that resulting from treatment with VP, namely, an increase in S256 and S269 phosphorylation and a decrease in S261 phosphorylation [17]. Using homology analysis of amino acid sequences of the AQP2 c-terminus, we identified ribosomal-S6-kinase (RSK) as a potential kinase that could also phosphorylate AQP2 at the S256 residue. This, along with the fact that RSK is downstream of EGFR signaling, prompted us to pre-treat our LLC-AQP2 cells with BI-D1870, an RSK inhibitor, prior to the addition of erlotinib. Using erlotinib alone as our positive control, we found that inhibition of RSK by BI-D1870 causes a significant reduction of erlotinib-induced phosphorylation at S256 (Fig. 1). Because we previously showed that erlotinib, similar to VP, induced AQP2 membrane accumulation, we next asked if BI-D1870 could decrease erlotinib-induced membrane accumulation.

**Figure 1.**
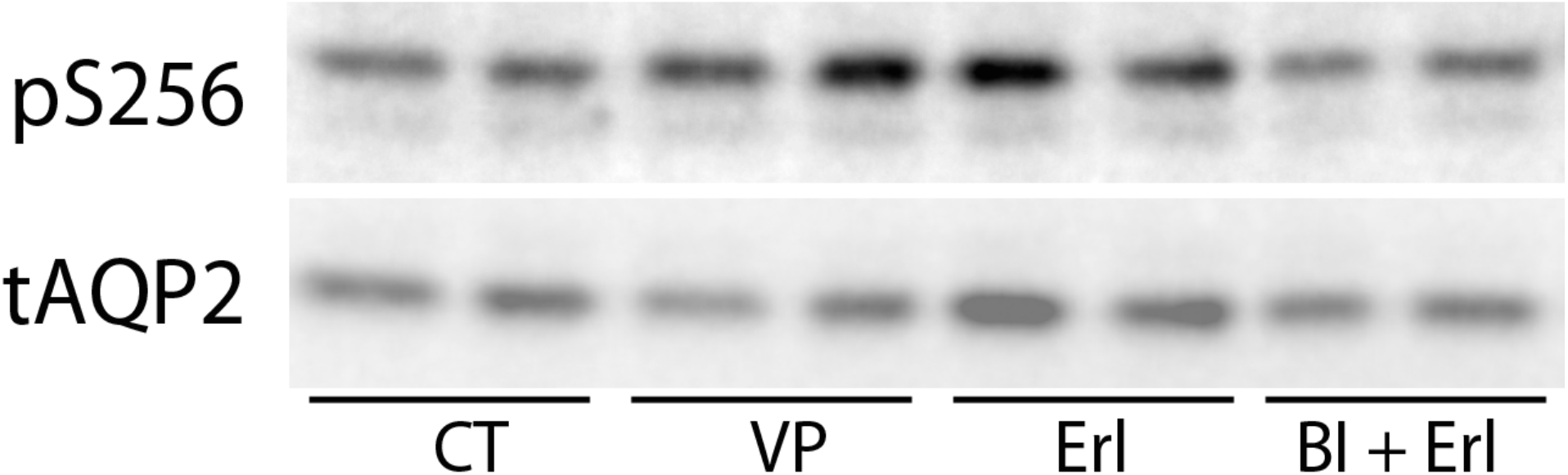
Inhibition of RSK decreases erlotinib-induced S256 phosphorylation of AQP2 in LLC-PK1 cells. Western blots were performed on lysates of LLC-PK1 cells stably expressing AQP2 and stained with specific phospho-AQP2 antibodies. Phosphoserine band intensities were quantified and normalized to their respective total AQP2 loading controls. Erlotinib increased S256 phosphorylation similar to VP, and this effect was significantly decreased in cells pre-treated with the RSK inhibitor BI-D1870. Data were analyzed using a Student t-test, and expressed as mean ± SD. (n=5)

### Inhibition of RSK decreases erlotinib-induced AQP2 membrane accumulation in cells

As shown in our previous work, erlotinib increases AQP2 membrane accumulation in our LLC-AQP2 cells, an effect also seen with VP, a positive control (Fig. 2, top panel). When pre-treated with the RSK inhibitor BI-D1870, erlotinib-induced membrane accumulation was greatly attenuated, with most AQP2 remaining in the cytoplasm as seen in untreated control cells (Fig. 2 bottom right). However, BI-D1870 did not affect vasopressin-induced membrane accumulation of AQP2 (Fig. 2, bottom middle), suggesting the effect is specific to the EGFR pathway and not the canonical PKA signaling pathway activated by VP. In order to confirm the effect of RSK inhibition on erlotinib-induced AQP2 membrane accumulation, we next used small interfering RNA (siRSK) to knockdown RSK in our cells.

**Figure 2.**
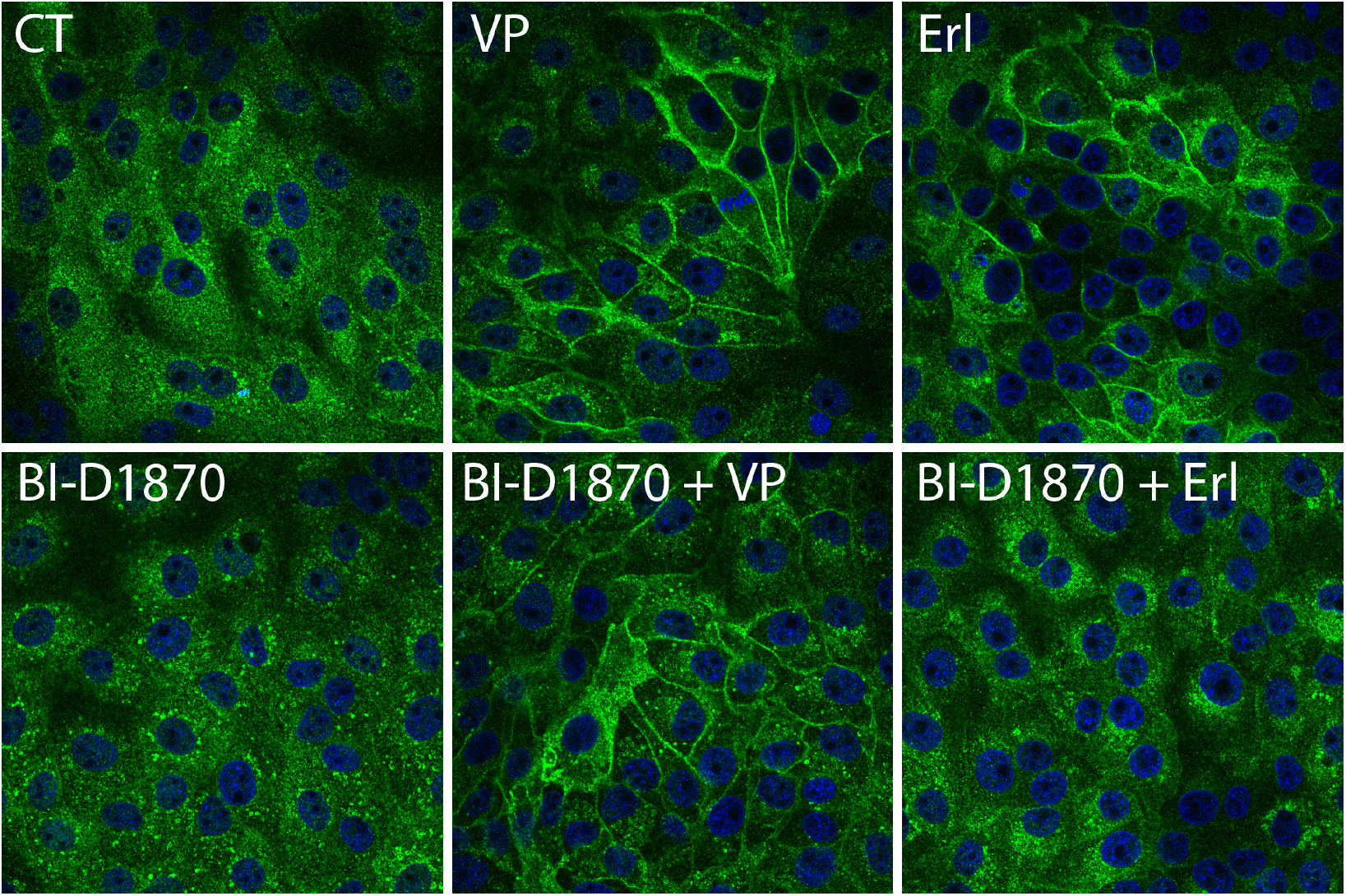
RSK inhibition abolished erlotinib-induced AQP2 membrane accumulation in LLC-AQP2 cells. Erlotinib-induced AQP2 membrane accumulation in LLC-PK1 cells is blocked by RSK inhibition. As expected, both VP and erlotinib (Erl) induced an increase in AQP2 membrane accumulation (top row). When pre-treated with RSK inhibitor, BI-D1870, erlotinib-induced AQP2 membrane accumulation is inhibited, and AQP2 is mostly in the cytoplasm (bottom right panel), similar to controls (CT). BI-D1870 does not inhibit VP-induced AQP2 membrane accumulation (bottom middle panel) and BI-D1870 alone does not cause visible changes in AQP2 localization (bottom right panel). These images are representative of 4 independent experiments (n=4)

### Knockdown of RSK decreases erlotinib-induced membrane accumulation of AQP2 and erlotinib-induced phosphorylation of AQP2 at S256

LLC-AQP2 cells were transfected with RSK siRNA (siRSK) for 48 hours and showed a significant decrease in RSK protein expression. Compared to control cells, those transfected with RSK siRNA showed significantly decreased S256 phosphorylation in response to erlotinib (Fig. 3A). Likewise, erlotinib treated cells transfected with siRSK demonstrated decreased apical membrane trafficking of AQP2 compared to cells transfected with scrambled siRNA (Fig. 3B, bottom right panel). While siRSK decreased erlotinib-induced ser256 phosphorylation and membrane trafficking of AQP2, it failed to decrease vasopressin-induced phosphorylation or membrane trafficking (Fig. 3B, bottom middle panel). This again suggests RSK as an effector in the EGFR pathway, and that this pathway is distinct from the canonical vasopressin-activated pathway.

**Figure 3.**
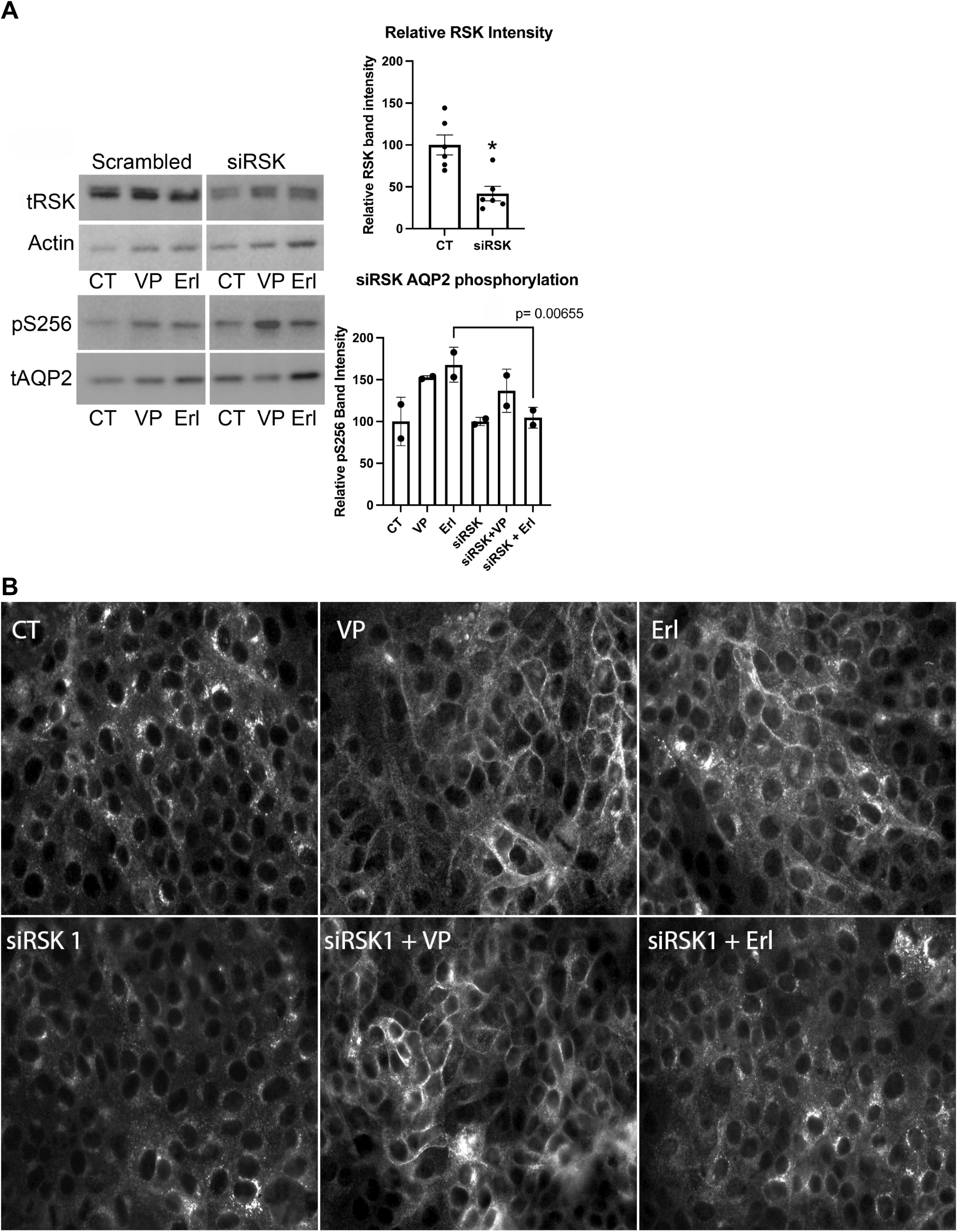
Downregulation of RSK with siRNA abolished erlotinib induced S256 AQP2 phosphorylation and AQP2 membrane accumulation. RSK protein abundance is significantly reduced after cells were incubated with siRNA specific for RSK protein (siRSK, top panel). When incubated with scrambled siRNA, increased phosphorylation of S256 is seen with VP and erlotinib treatment (bottom). The phosphorylation of S256 AQP2 induced by erlotinib is significantly reduced in cells incubated with siRSK, but not in VP-treated cells. This is representative of 5 independent experiments (n=5). After incubating cells with scrambled or siRSK for 48 hours, cells were exposed to VP or erlotinib. In cells incubated with scrambled siRNA (top row), VP and erlotinib increased AQP2 membrane accumulation. In cells incubated with siRSK, VP continued to induce AQP2 membrane accumulation, but erlotinib no longer induced membrane accumulation of AQP2. These images are representative of 3 independent experiments (n = 3).

### RSK is expressed in principal cells of rat kidney collecting ducts along with AQP2

Next, we sought to confirm the presence of RSK in kidney tissue of our rodent model. We found that there is a differential expression of RSK (Fig. 4, green) in the kidneys, with a predominance of RSK in the collecting duct principal cells of renal medulla and papilla, as identified by their staining for AQP2 (Fig. 4, red). The expression of RSK in the cortical collecting duct was lower than in the medulla.

**Figure 4.**
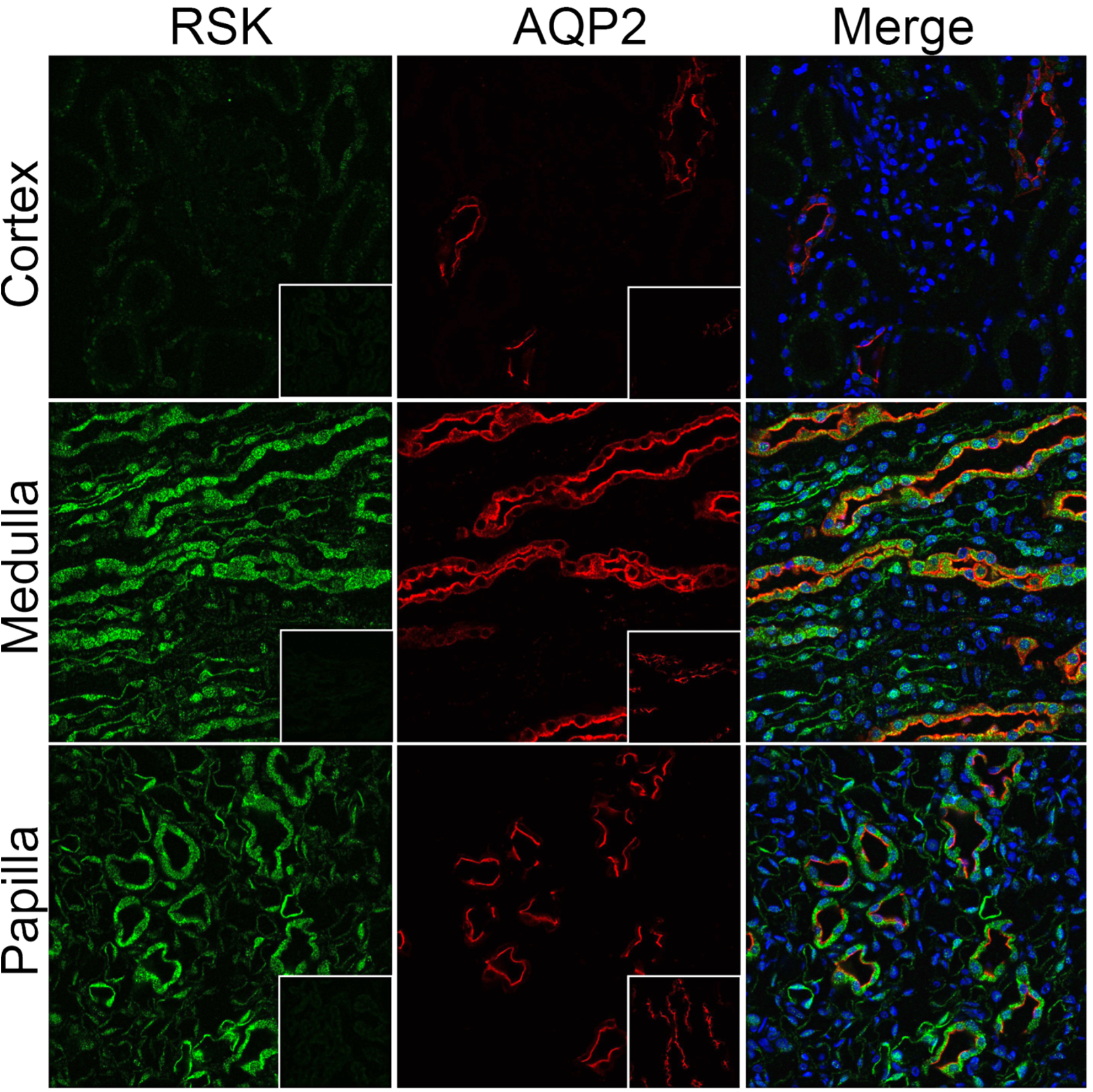
There is predominance of RSK in the kidney principal cells and co-localizes with AQP2 in the renal medulla and papilla. Immunohistochemistry shows RSK (green, left column) and AQP2 (red, middle column) in cortex (top row), medulla (middle row) and papilla (bottom row). Merged panels are shown in the right column. RSK was heavily expressed in principal cells in the collecting ducts in the medulla and papilla, co-localized with AQP2 in red. Expression of RSK is to a lesser degree in the cortex. The insets in the left column show the lack of specific RSK staining in the presence of a blocking peptide. The insets in the middle column show the expected AQP2 (red) staining in tissues double stained after application of RSK blocking peptide. These images are a representative of three independent experiments on three different animals. (n=3)

### Erlotinib-induced AQP2 membrane accumulation is decreased by inhibition of RSK in kidney principal cells

Since RSK is clearly expressed in collecting duct principal cells, we pre-treated kidney slices with the RSK inhibitor BI-D1870 to determine whether it also inhibited erlotinib-induced AQP2 membrane accumulation in situ. Similar to our finding in cultured cells, erlotinib was no longer able to induce AQP2 membrane accumulation in kidney slices pre-treated with BI-D1870 (Fig. 5, bottom right panel). These encouraging results further suggested a physiological role of RSK in AQP2 trafficking. As homology analysis of the amino acid sequence in AQP2’s c-terminus predicted that RSK may phosphorylate AQP2 at S256, and because the RSK inhibitor BI-D1870 reduced AQP2 S256 phosphorylation, we next asked whether RSK is capable of directly phosphorylating AQP2.

**Figure 5.**
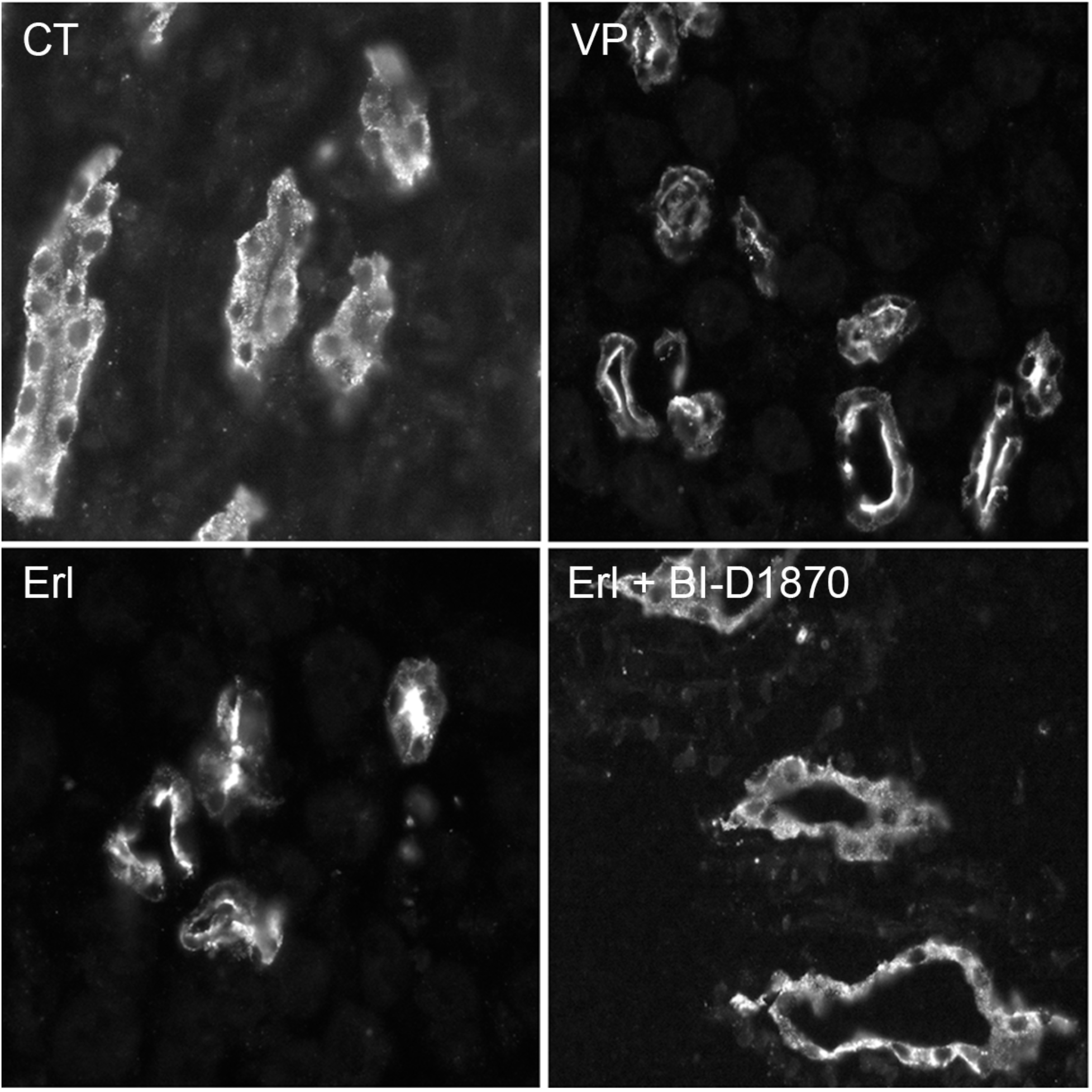
Inhibition of RSK decreases erlotinib-induced membrane accumulation of AQP2 in rat kidneys. Kidney slices were pre-treated with the RSK inhibitor, BI-D1870, followed by erlotinib treatment for 30 min. Apical membrane accumulation of AQP2 is noted in kidney slices treated with VP and erlotinib (Erl), but pre-treatment with BI-D1870 abolishes Erl-induced apical membrane accumulation as AQP2 instead remains diffusely distributed in the cytoplasm of principal cells. These images are representative of 4 independent experiments from 4 different animals (n=4)

### RSK directly phosphorylates AQP2 at ser256 *in vitro*

We tested if RSK was able to directly phosphorylate AQP2 *in vitro*. A recombinant c-tail of AQP2 was incubated with either PKA, our positive control, or RSK, and radioisotope phosphorus ^32^P. In our positive control, we confirmed that PKA can directly phosphorylate the c-tail ofAQP2, and as expected this phosphorylation was inhibited by the known PKA inhibitor H89. When incubated with a mutated c-tail of AQP2 in which the S256 was mutated to unphosphorylatable alanine (S256A), PKA no longer caused AQP2 c-tail phosphorylation (Fig. 6 left). We performed the same assay using RSK and similarly showed that RSK can directly phosphorylate AQP2 c-tail but cannot when serine is replaced by alanine. When the RSK inhibitor BI-D1870 was added, RSK was no longer able to phosphorylate AQP2 in its c-terminus (Fig. 6, right). This finding confirms that RSK can directly phosphorylate AQP2 at S256, and may, therefore, be the terminal kinase in the erlotinib-activated pathway of AQP2 phosphorylation and membrane accumulation.

**Figure 6.**
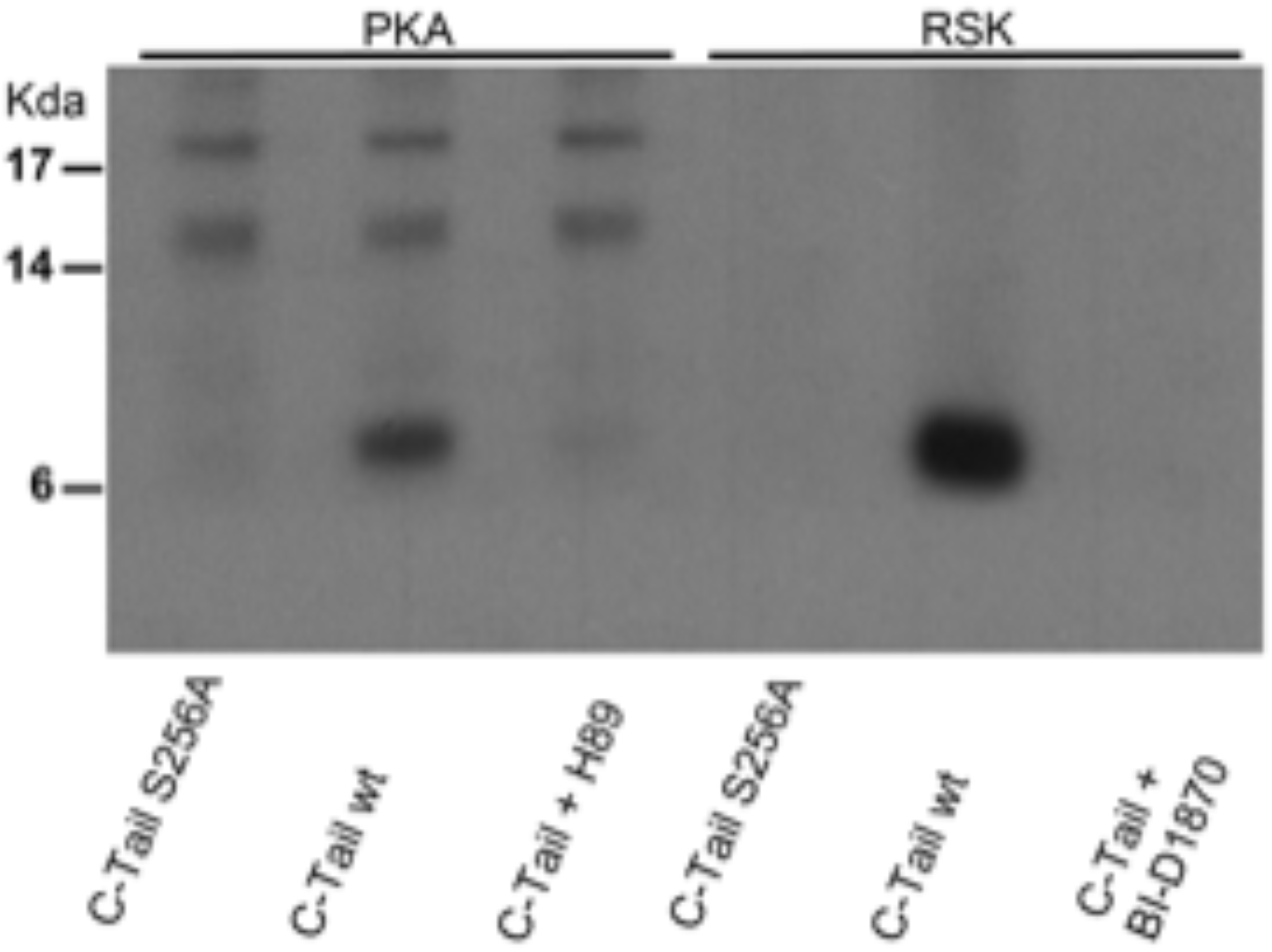
RSK can directly phosphorylate AQP2 at S256. A C-terminal tail (∼7kDa) construct of AQP2 was incubated with RSK and radioisotope ^32^P *in vitro*. As a positive control, the AQP2 c-tail was incubated with recombinant protein kinase A (PKA), and as expected our assay demonstrated phosphorylation of the AQP2 c-tail. A serine to alanine mutation behaves functionally as a permanently dephosphorylated residue, and recombinant AQP2 c-tail with this S256A mutation failed to show phosphorylation by PKA. H89, a non-specific PKA inhibitor, prevents the ability of PKA to phosphorylate AQP2 c-tail (left three lanes). RSK (right three lanes) can also directly phosphorylate the AQP2 c-tail, similar to PKA, and cannot phosphorylate the c-tail S256A mutation. BI-D1870 inhibits the ability of RSK to phosphorylate AQP2 c-tail. This image is a representative of three independent experiments (n=3).

### Upregulation of phosphoinositide-dependent kinase 1 (PDK1), an activator of RSK, increases AQP2 membrane trafficking

As we demonstrated that RSK can directly phosphorylate AQP2 and inhibition of RSK abolished erlotinib-induced AQP2 membrane accumulation, we next evaluated whether direct activation of RSK could cause AQP2 membrane accumulation. Phosphoinositide-dependent kinase 1 (PDK1) is a kinase recruited by RSK as part of a sequence of events leading to its activation [27]. PDK1 phosphorylates the S221 residue on the N-terminus of RSK, a phosphorylation state thought to be integral to RSK’s ability to phosphorylate substrates. Therefore, we used PS210, a PDK1 activator, to see if we could affect AQP2 trafficking by activating RSK at its S221 residue.

Here, we found that treating cells with PS210 caused AQP2 accumulation in the plasma membranes of LLC-AQP2 cells, similar to erlotinib or VP treated cells (Figure 7). Using GSK2334470, an inhibitor of PDK1, we found that erlotinib or PS210-induced AQP2 membrane accumulation was inhibited. As shown in our previous work, the non-specific kinase inhibitor H89 decreased both vasopressin and erlotinib-mediated AQP2 membrane trafficking [17]. PS210-mediated membrane trafficking was also decreased by H89. These data suggest that PDK1, by way of RSK, is part of the signaling pathway of AQP2 membrane trafficking induced by the EGFR inhibitor erlotinib.

**Figure 7.**
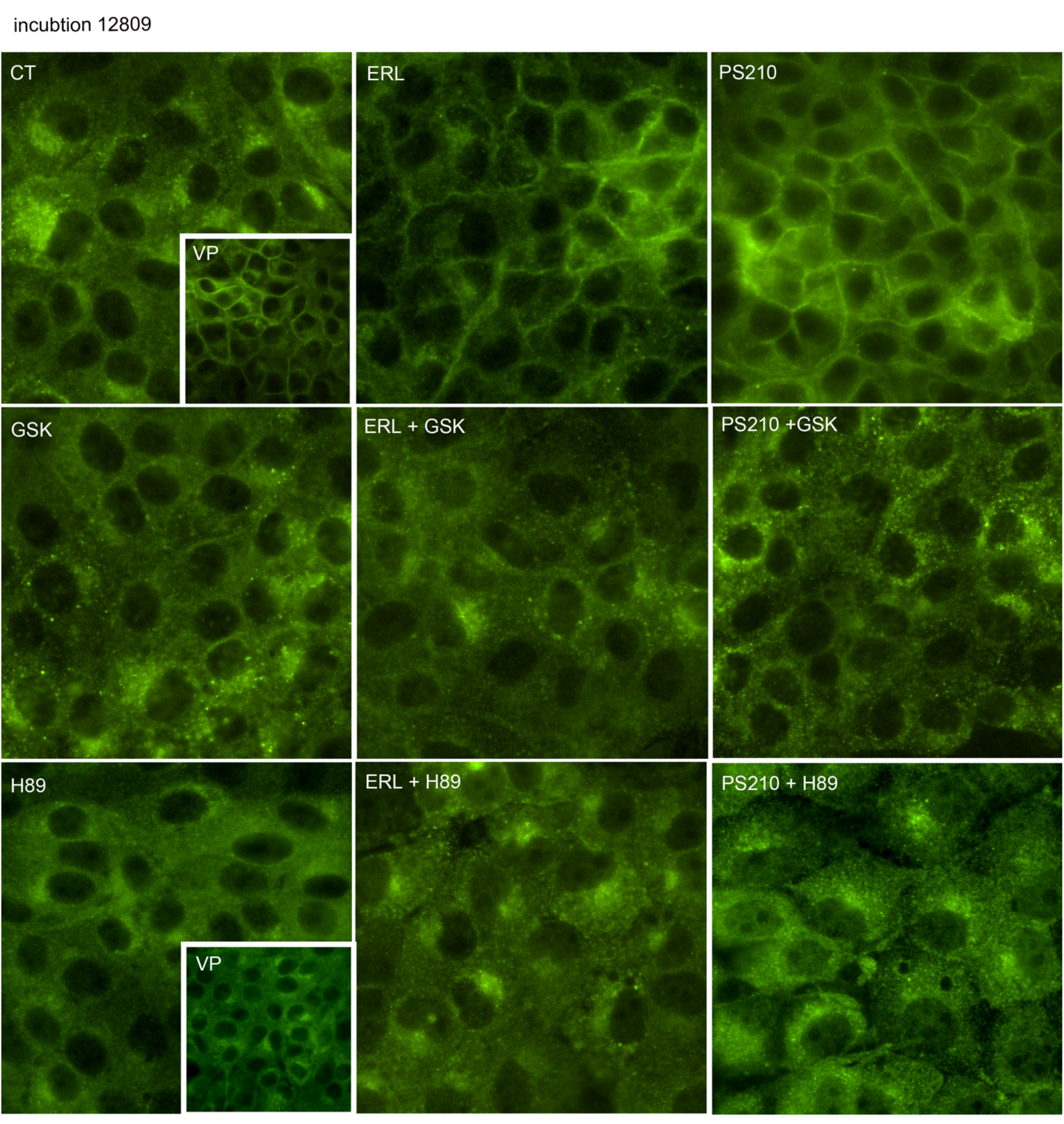
Phosphoinositide-dependent kinase 1 (PDK1) activation increases AQP2 membrane accumulation. PS210, a PDK1 activator, caused AQP2 membrane accumulation as also seen with erlotinib or vasopressin (top row). Membrane accumulation of AQP2 induced by either erlotinib or PS210 is decreased by the PDK1 inhibitor GSK-470 (middle row). The non-specific kinase inhibitor H89 decreased membrane trafficking induced by PS210 in addition to membrane trafficking in vasopressin and erlotinib treated cells (bottom row).

### Erlotinib fails to induce AQP2 S256 phosphorylation in RSK knockout cells

Next, we tested whether erlotinib was able to cause AQP2 phosphorylation in the absence of RSK. Using CRISPR techniques, we created an LLC-AQP2 RSK knockout cell line (Fig 8). Consistent with our previous studies, both erlotinib and VP increased S256 phosphorylation in our regular LLC-AQP2 cells (Fig 8). RSK KO cells failed to show an increase in AQP2 S56 when treated with erlotinib, but did show an increase in S256 phosphorylation when treated with VP. This demonstrates that RSK is indispensable to erlotinib-induced AQP2 signaling pathway, and confirms that this pathway is separate from that initiated by VP.

**Figure 8.**
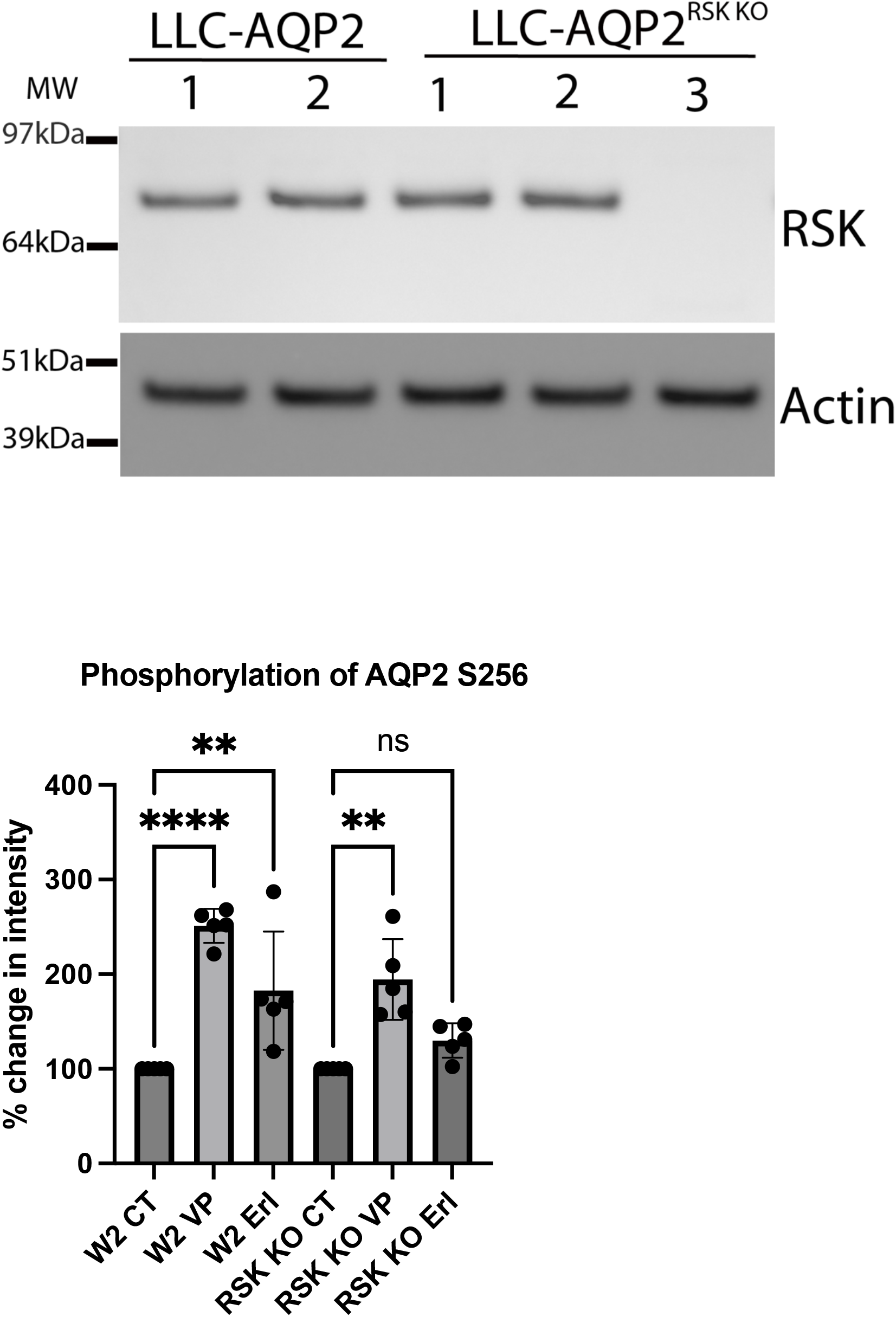
RSK knockout cells do not show increased AQP2 S256 phosphorylation in response to erlotinib. An RSK knockout in the LLC-AQP2 cell line (RSK KO) was created using CRISPR/Cas9 and compared to regular LLC-AQP2 cells (W2) with intact RSK. Knockout was verified by western blot with clone 3 showing successful knockout. Cells were treated with erlotinib (20μM), or VP (10nM) for 30 or 15 min respectively. Western blots were performed on lysates of cells and stained with specific phospho-AQP2 antibodies. Both cell lines showed increased AQP2 phosphorylation in response to the positive control vasopressin, while only the cells with intact RSK demonstrated increased AQP2 S256 in response to erlotinib. This result was repeated 5 times in duplicate. Mean ± SD, N=5, \*\**P*<.01, \*\*\*\**P*<.0001).

## Discussion

In this study, we found that erlotinib-induced AQP2 membrane accumulation relies on activation of RSK, and that activation of RSK with the PDK1 activator PS210 increases AQP2 phosphorylation and membrane accumulation. Our results identify RSK as the most probable terminal kinase functioning in the novel AQP2 signaling pathway induced upon EGFR inhibition with erlotinib.

RSK’s structure is unique and features two distinct kinase domains connected by a linker region. The C-terminal kinase domain (CTKD) is related to the CAMK family while the N-terminal kinase domain (NTKD) is related to the family of protein kinases A, G and C. It is thought that two phylogenetically distinct kinases merged in the course of evolution to create RSK [27]. RSK’s activation is complex. ERK binds to the CTKD resulting in phosphorylation of T573 of the CTKD. The CTKD then in turn phosphorylates S380 in the linker region. The phosphorylated residue S380 serves as a docking site for another kinase called phosphoinositide-dependent kinase-1 (PDK1), which phosphorylates S221 in the NTKD resulting in RSK activation. The NTKD is thought to phosphorylate RSK’s known substrates, and phosphorylation of the S221 residue in the NTKD is considered necessary for RSK to act on its substrates. Even though this activation cascade is considered canonical, more recent studies have shown that RSK activation can be modulated independently of the CTKD, and NTKD activation alone is sufficient for RSK to induce phosphorylation of its substrates [37].

One conceptual issue in ascribing this role to RSK is the apparent incongruency that erlotinib, an inhibitor of the EGFR, is able to *increase* the activity of one of EGFR’s downstream effectors in the MAPK pathway, namely RSK. If erlotinib is known to inhibit MAPK activity, how then is erlotinib able to activate RSK? One possible explanation may relate to the spatiotemporal localization of RSK’s activator PDK1. EGF is known to cause PDK1 to localize to the plasma membrane [38]. As RSK is often activated in the cytosol [26], EGFR inhibition with erlotinib may cause increased PDK1 accumulation in the cytosol as opposed to the membrane, and thereby increase RSK activity. Whether this is the mechanism by which erlotinib activates RSK, or whether EGFR inhibition instead perturbs a negative feedback loop tonically inhibiting RSK, remains to be seen and will be the subject of future studies.

The mechanism of EGFR activation by its ligands is complex and remains a matter of debate. In an inactive state the EGFR exists as monomers in the cell membrane. Upon ligand binding, these EGFR monomers dimerize to form either homodimers or heterodimers with other members of the Erb-B family (eg. HER2, HER3, or HER4). Dimerization brings the intracellular C-terminal tyrosine kinase domains within proximity of each other resulting in autophosphorylation and activation of several downstream pathways. Following activation, the EGFR is internalized and trafficked to the early endosomal compartment [23, 25]. The rates of internalization as well as the downstream signaling effects of the EGFR depend on both the type of ligand binding the EGFR as well as the composition of the heterodimer [25, 39]. Interestingly, it has recently been shown that certain classes of EGFR inhibitors induce dimerization and internalization, while others do not [40]. Furthermore, it appears that internalization, dimerization, and ERK phosphorylation all vary according to the class of EGFR inhibitor used [40]. Group 1 tyrosine kinase inhibitors, of which erlotinib is a member, are among those that cause dimerization and internalization. It is currently unknown what effects, if any, dimerization or internalization of inhibited EGFR have on downstream EGFR signaling. Studies examining whether other classes of EGFR inhibitors share erlotinib’s effect on AQP2 and RSK will help illuminate whether these mechanisms of inhibition are relevant to EGFR-mediated AQP2 signaling.

The opposing effects of VP and EGF on water reabsorption in the kidney has long been suspected [18], and studies confirming that EGF induces a water diuresis in sheep and rats were published in the 1990s [19, 20]. Our group previously showed that EGF attenuated AQP2 S256 phosphorylation and EGFR inhibition with erlotinib increased AQP2 S256 phosphorylation, AQP2 membrane accumulation, and urine osmolarity in mice with lithium-induced NDI [17].

Literature in recent years has found an emerging role for EGF in several disparate fields of renal physiology. In addition to water homeostasis, EGF regulates magnesium and sodium handling [41–43], and has also been shown to play a role in repair of AKI and progression of CKD [44–47]. mRNA levels of EGF are higher in the kidney than any other human organ tissue by over 100-fold [46]. While tissue protein levels of EGF in the kidney are also higher than most other tissues, it appears the vast majority of EGF generated in the kidney is secreted in urine [48, 49]. It is also clear that urinary EGF is produced in the kidney rather than filtered from the systemic circulation [50, 51]. During AKI, there is increased production of EGFR ligands and EGFR signaling [52]. Studies of chronic kidney disease show that urinary EGF (uEGF) declines with progression of renal fibrosis, and uEGF has been shown to be highly predictive of CKD progression [45–47, 53].

In this way, a picture emerges of urinary EGF as a paracrine signaling molecule acting at various sites along the nephron. Throughout the lifespan of the kidneys, EGF participates in tissue repair and remodeling after episodes of acute kidney injury. Over time, uEGF levels fall as progression of renal fibrosis is accompanied by a decline of EGF production. At the same time, uEGF regulates water and electrolyte homeostasis in the distal nephron. EGF’s dual role of regulating tissue repair and water homeostasis may account for the water diuresis often observed in the recovery phase of acute tubular necrosis.

The discovery of detailed signaling mechanisms in the novel EGFR-AQP2 pathway promises new therapeutic targets that may be exploited for treatment in the field of water balance disorders where many unmet clinical needs remain. Nephrogenic diabetes insipidus (NDI) is characterized by excessively dilute urine resulting from an inability to traffic AQP2 to the membrane of the collecting duct in response to increased circulating vasopressin [14]. A lack of effective treatments for NDI often results in poor quality of life with some patients drinking up to 20 liters daily to replenish their urinary losses [16]. Our previous study showed erlotinib was able to partially reverse NDI in mice [17]. While erlotinib is often useful for cancer therapy, it would be impractical to treat NDI patients with EGFR inhibitors in light of their broad biological activity which includes the side effect of severe diarrhea. Therefore, the identification of more specific mediators such as RSK and its regulatory proteins such as PDK1 may offer a novel therapeutic target more appropriate for patients with water balance disorders.

While the pathology of NDI centers around a lack of AQP2-mediated water resorption, water balance disorders characterized by excessive water reabsorption are far more commonly encountered in clinical practice. This group includes the hypervolemic states of cirrhosis and congestive heart failure, as well as the syndrome of inappropriate anti-diuretic hormone (SIADH). A class of medications known as “vaptans” reduces VP activity by blocking the V2R. Vaptans have been shown to decrease edema, aid weight loss, and improve quality of life in patients with diuretic-resistant cirrhosis and congestive heart failure [6–9, 12]. One study even showed long term vaptan use was associated with improved mortality in cirrhotics with hyponatremia [10]. Despite these promising studies, vaptans are not regularly used in these populations due to many adverse effects which include rapid overcorrection of sodium, hepatotoxicity, and progression of kidney disease [11–13]. The novel EGFR-AQP2 pathway may provide alternative pharmacologic targets to inhibit VP activity, allowing treatment of these diseases without the drawbacks accompanying vaptans. It should be noted here that other non-canonical pathways of AQP2 membrane accumulation have been reported [54–57], yet it remains to be seen whether any of them will yield a clinically useful target for treatment of water balance disorders.

Taken together, our results identify RSK as the kinase phosphorylating AQP2 in the novel signaling pathway induced by erlotinib. Our inquiries have led to the discovery of a new agent able to effect AQP2 phosphorylation and membrane trafficking: the PDK1 activator PS210. The potential to find new therapeutic targets for the treatment of water balance disorders warrants further study of this non-canonical signaling pathway on a molecular level.

## Acknowledgements

This work was supported by the National Institutes of Health (NIH) grant DK096586 (D. Brown). P. W. Cheung was supported was supported by NIH K-award DK115901, and a generous philanthropic gift from Donald Glazer. C.C.S.L. was supported by a NIH training grant 5T32DK007540. The Nikon A1R and the Zeiss LSM 800 with Airyscan confocal in the Program in Membrane Biology (PMB) Microscopy Core were purchased using NIH Shared Instrumentation Grants 1S10RR031563-01 and 1S10OD021577-01 (DB), respectively. Additional support for the Program in Membrane Biology Microscopy Core came from the Boston Area Diabetes and Endocrinology Research Center (DK057521) and the Massachusetts General Hospital (MGH) Center for the Study of Inflammatory Bowel Disease (DK043351).

## Notes

### Competing Interest Statement

The authors have declared no competing interest.

